# Identifying the engagement of a brain network during a targeted tDCS-fMRI experiment using a machine learning approach

**DOI:** 10.1101/2022.09.12.507591

**Authors:** A. B. Shinde, S. Mohapatra, G. Schlaug

## Abstract

Transcranial direct current stimulation (tDCS) can noninvasively modulate behavior, cognition, and physiologic brain functions depending on polarity and dose of stimulation as well as montage of electrodes. Concurrent tDCS-fMRI presents a novel way to explore the parameter space of non-invasive brain stimulation and to inform the experimenter as well as the participant if a targeted brain region or a network of spatially separate brain regions has been engaged and modulated. We compared a multi-electrode (ME) with a single electrode (SE) montage and both active conditions with a no-stimulation (NS) control condition to assess the engagement of a brain network and the ability of different electrode montages to modulate network activity. The multi-electrode montage targeted nodal regions of the right Arcuate Fasciculus Network (AFN) with anodal electrodes placed over the skull position of the posterior superior temporal/middle temporal gyrus (STG/MTG), supramarginal gyrus (SMG), posterior inferior frontal gyrus (IFG) and a return cathodal electrode over the left supraorbital region. In comparison, the single electrode montage used only one anodal electrode over a nodal brain region of the AFN, but varied the location between STG/MTG, SMG, and posterior IFG for different participants. Whole-brain rs-fMRI was obtained every three seconds. The tDCS-stimulator was turned on at 3 minutes after the scanning started. A 4D rs-fMRI data set was converted to dynamic functional connectivity (DFC) matrices using a set of ROI pairs belonging to the AFN as well as other unrelated brain networks. In this study, we evaluated the performance of five algorithms to classify the DFC matrices from the three conditions (ME, SE, NS) into three different categories. The highest accuracy of 0.92 was obtained for the classification of the ME condition using the K Nearest Neighbor (KNN) algorithm. In other words, applying the classification algorithm allowed us to identify the engagement of the AFN and the ME condition was the best montage to achieve such an engagement. The top 5 ROI pairs that made a major contribution to the classification of participant’s rs-fMRI data were identified using model performance parameters; ROI pairs were mainly located within the right AFN. This proof-of-concept study using a classification algorithm approach can be expanded to create a real-time feedback system at a participant level to detect the engagement and modulation of a brain network that spans multiple brain lobes.

## Introduction

Experimental studies revealed evidence that transcranial direct current stimulation (tDCS) modulates brain activity and affects behavior, cognition, and sensorimotor skills among others when that activity draws on targeted brain regions or entire brain networks [1–4]. tDCS applies a constant electric current to targeted brain regions through their scalp access points typically using one or more anodal electrodes and one or more cathodal electrodes [5,6]. The current induces subthreshold changes across the neuronal membrane potentially modulating the neuronal firing depending on the polarity and dose of the current as well as the montage of electrodes. In general, brain regions targeted with anodal tDCS shows increased excitability while cathodal tDCS decreases it, as commonly shown in the studies stimulating motor cortex [3,7–9]. Moreover, modulatory effects of tDCS have been shown in non-motor brain regions as well [10–13].

There is compelling evidence that targeted tDCS can not only modulate local intrinsic brain activity or the activity directly under the targeted brain region, but also influence activity in connected brain regions spanning longer distances in the brain. Modulation of multiple connected regions or entire networks may account for the effects on complex behavior and cognition seen in neurologic and psychiatric disorders [3,8,14–22]. One network that is uniquely situated, spans multiple lobes, and could be targeted by identifying nodal cortical access points, is the Arcuate Fasciculus Network (AFN). The AFN includes brain regions in the posterior superior and middle temporal gyri, the inferior parietal lobule (supramarginal gyrus), and the inferior frontal gyrus and connections with several regions in between such as the inferior sensorimotor cortex. This network is ideal for examining whether targeting particular cortical access points with different electric montages can differentially modulate rs-fMRI using concurrent tDCS-fMRI experiments.

Concurrent tDCS-fMRI has been used to reveal neural correlates of stimulation using various MR acquisition methods including resting-state fMRI, and varying dose and montage to test whether or not targeted brain regions can be engaged, and their connections be modulated [5,23]. Machine learning methods have been explored to characterize rs-fMRI, often grouped in two types: unsupervised and supervised [24]. Unsupervised methods focus on understanding healthy brain and its dynamics such as matrix decomposition and clustering to identify brain functional networks [25,26]. On the other hand, supervised learning methods use rs-fMRI data to classify ‘patient vs controls’ or to predict disease prognosis [27–29]. Despite widespread use of machine learning methods for rs-fMRI classification, the use of machine learning techniques in tDCS-fMRI studies is limited and has been restricted to binary classification questions [30–34]

In this paper we describe a supervised learning approach to evaluate the engagement of a targeted brain network, the Arcuate Fasciculus Network (AFN) using a tDCS-fMRI experiment. In the supervised learning approach, we classify the rs-fMRI data into three groups based on stimulation condition and electrode montage (Multielectrode vs Single Electrode and both vs a No-Stimulation condition). This is followed by identifying brain regions that show strong differential effects of tDCS depending on electrode montage.

## Methods

### Participants

A total of thirty-three (33) participants were recruited in the greater Boston area as well as in the greater Amherst area (Females= 14; mean age=36.7). Participants (n = 12) in the greater Boston area were scanned on a 3T GE scanner at Beth Israel Deaconess Medical Center (BIDMC) while participants recruited in the greater Amherst/Pioneer Valley region (n = 21) were scanned on a 3T Siemens wide bore scanner. All participants were right-handed according to the Edinburgh Handedness Inventory (Oldfield, 1971), had no history of neurologic or psychiatric conditions, and had no contraindications to undergo MRI or tDCS as verified by a safety checklist. The Institutional Review Boards of Beth Israel Deaconess Medical Center and University of Massachusetts Amherst approved this study, and all participants gave written informed consent. All participants underwent one mock session prior to the concurrent tDCS-fMRI sessions in which they were familiarized with the setup and were given a test stimulation of the 4mA stimulation to make sure that they were able to tolerate the intensity of stimulation and were comfortable in the MR scanner environment. Participants were then assigned to one of three concurrent tDCS-fMRI sessions on their first day of the experiment typically alternating between one of the stimulation montages and the no-stimulation session starting first; this was followed on subsequent days by either one of the remaining two sessions. Not all participants were able to participate in all sessions. To ensure that there was not a systematic difference between the BIDMC and UMass MRI sessions due to factors not controlled for, an initial analysis compared the BIDMC no-stimulation (NS) sessions (n = 5) with the first 5 UMass NS sessions. An ‘interhemispheric functional connectivity measure’ was calculated as the simple average of all possible interhemispheric functional connectivities between homotopic ROIs on either hemisphere from every session, and a t-test was conducted. ROIs used for this calculation are defined in the section ‘Selection of ROI pairs.’ This test did not show a significant difference between sites, with p>0.05. Therefore, the BIDMC and UMass data were used as a single dataset for all further analyses.

### Electrode placement

Prior to a participant entering the MRI room, we placed MR compatible rubber electrodes on each participant’s scalp using the 10-20 Electroencephalogram (EEG) system as a guide for the initial identification of scalp targets. The electrode targets were selected to stimulate nodal cortical regions of the AFN. In the multielectrode (ME) condition, we placed three round anodal electrodes (diameter of 3cm each) over the scalp position of the supramarginal gyrus (SMG), the posterior inferior frontal gyrus (IFG) and the posterior STG/MTG junctional region. The IFG electrode location was placed halfway between C6 and F8, the SMG electrode location was placed one third of the distance between C6 and CP4, and the STG/MTG electrode location was placed halfway between CP6 and TP8. A round cathodal electrode (diameter of 5cm) was placed over the contralateral supra-orbital region (approximately corresponding to FP1). In the single electrode (SE) condition, we placed a round electrode (diameter of 4cm) over either the SMG, the pIFG, or the posterior STG/MTG junction and the cathodal electrode again over the left fronto-orbital region (FP1). The electrode size and the currents were selected to have similar maximum charge density values (ME: 0.2132 C/cm^2^ and SE: 0.2292 C/cm^2^). After cleaning the targeted scalp locations with alcohol, rubber electrodes lathered with approximately 2mm thick layer of Ten20 neurodiagnostic electrode paste (Weaver and company, Aurora, CO, USA) were placed. Electrodes were held in place using a self-adhesive bandage and hypoallergenic medical tape. Electrode connectors were adjusted to avoid any crossover between wires. The no-stimulation (NS) data was acquired while participants had either the single or multi-electrode stimulation montage placed over their head and were informed that stimulation might be applied but was not actually applied in that session. Seven participants out of 33 participants from the no-stimulation (NS) group, participated in an MRI session where no electrodes were placed over the participant’s scalp; these participants were naive to the stimulation at the time. The NS condition was a control condition without any stimulation, not even a sham stimulation. Even when electrodes were mounted, we did not ramp up/down the stimulation as is typically done in a sham stimulation condition, since this effect could potentially engage the network that we were trying to identify with either the ME and SE conditions in comparion to the NS condition.

### Concurrent tDCS-fMRI

After placing electrodes on the scalp, participants were placed inside the MRI bore in a supine position. Electrodes were connected to the MR-safe connector box. The connector box was connected to the RF filter panel inside the MR room using the MR-safe connector cable. A second connector cable was used in the MR control room connecting the RF filter panel to the MR conditional connector box. The MR conditional connector box converted the stimulation signal generated using Neuroconn DCMC stimulator into a signal that was carried through a connector cable. All the aforementioned components were provided with the Neuroconn DCMC stimulator device (Neurocare Group, Germany). A schematic diagram of these connections is shown in figure 1a.

**Figure 1:**
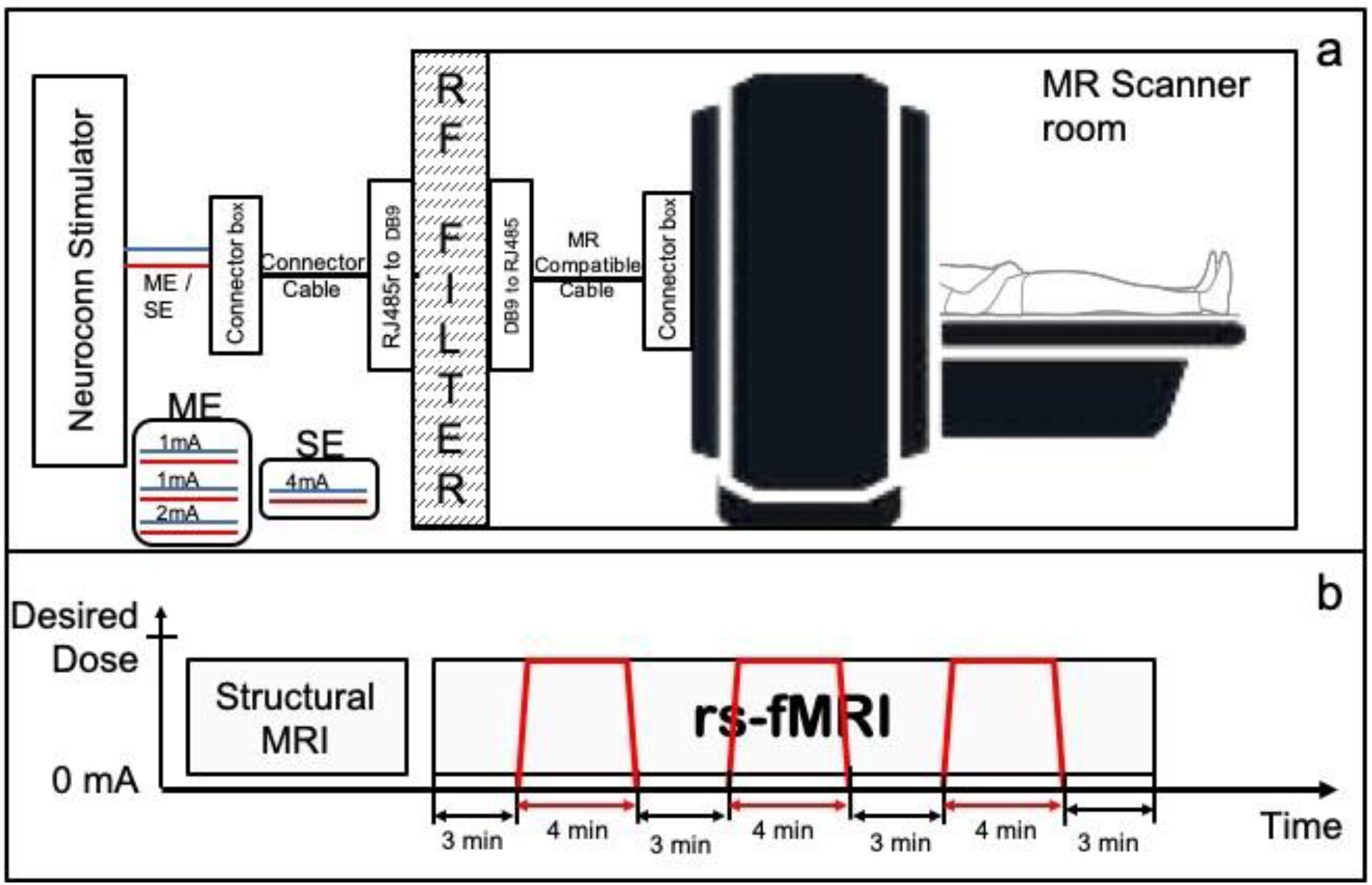
a) Concurrent tDCS-fMRI setup. b) Timing diagram for MRI experiment. Red colored lines show the application of tDCS during the 24-minute long rs-fMRI scan.

**Figure 2:**
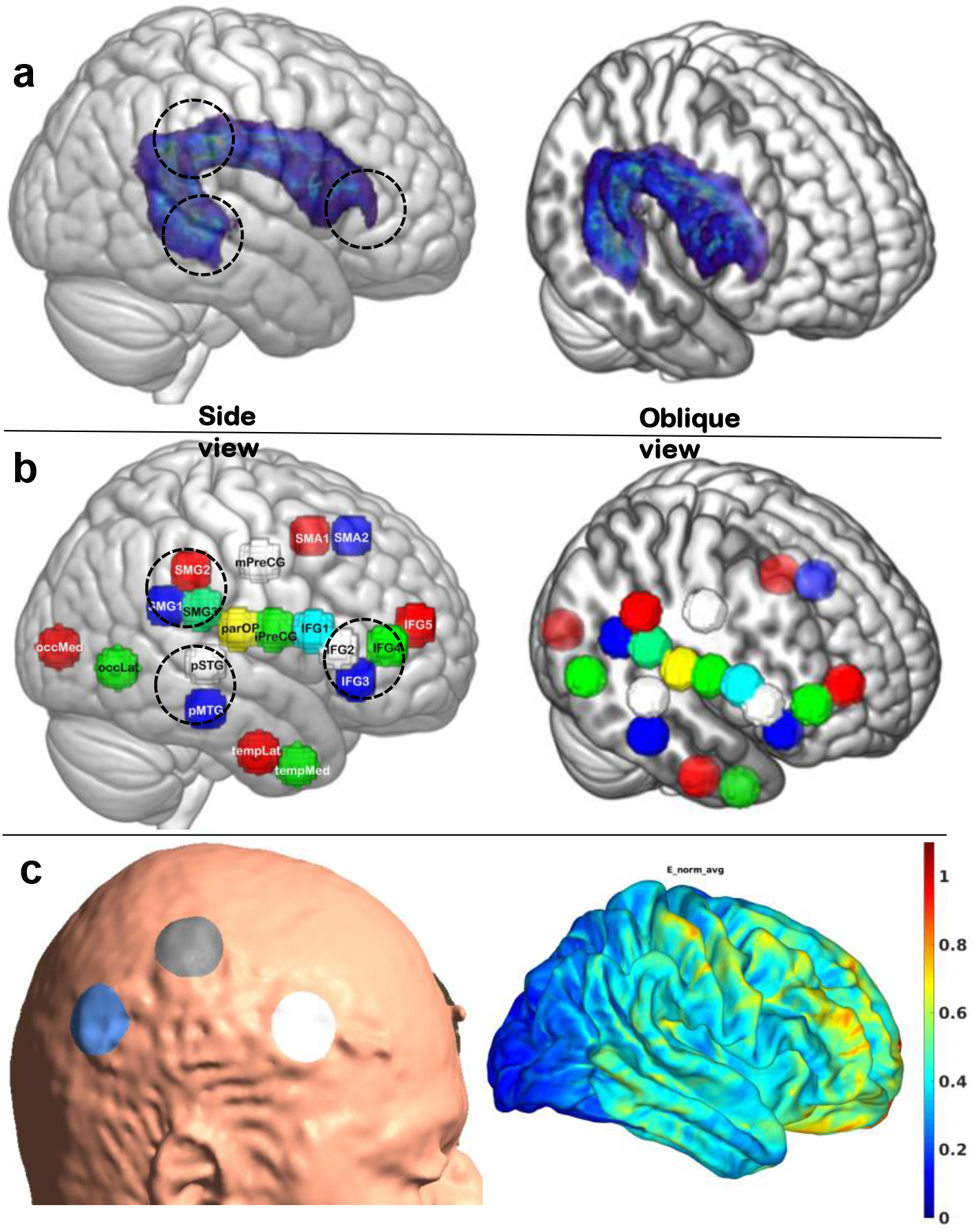
(a) Representative image of the right hemispheric Arcuate Fasciculus tract shown in standard brain and the canonical location of three nodal regions of the AFN targeted by brain-stimulation (dashed circles in 2a). tDCS electrodes were placed on the scalp directly above the brain regions marked with circles. (b) Anatomically identified spherical ROIs representing cortical regions within and outside of the AFN (shown here for the right hemisphere, but mirror left hemisphere regions were identified as well). Side and oblique views are shown to demonstrate the mesial location of AF tract and ROIs. (c) Shows electrode placement for ME montage for one of the participant and Electric field distributions simulated using SimNIBS software [39].

After a participant was situated in the scanner and all connections were secured, a T1-weighted 3D MPRAGE sequence (resolution= 1.0 × 1.0 × 2.0*mm*^3^; TR/TE 1490/3.36ms; flip angle = 9°; matrix = 256 × 256; field of view = 256 × 256*mm*^2^) with sagittal acquisition and 2mm slice thickness was obtained to speed up the acquisition of an anatomical image. This 3D anatomical image set was used to confirm the accuracy of the electrode location with underlying brain regions that we intended to target. If the electrodes were placed inaccurately (meaning that they did not overlay the target region), then the participant was moved out of the MR bore, the electrode position was adjusted, and the corrected placement was again confirmed with a fast T1-weighted acquisition. When electrodes were accurately placed, a 24-min rs-fMRI scan using a gradient-echo echo planar imaging sequence (TR of 3s, TE of 31.0ms; flip angle of 90 degrees; field of view = 210×210*mm*^*2*^; and voxel size of 2.5 × 2.5 × 2.5*mm*^3^) was obtained. The stimulator was turned ON and OFF during this 24-minute scan with ON periods being 4-minutes long (with a 30 second ramp-up and ramp-down) and OFF periods 3-minutes long, flanking the ON-stimulation at the beginning and the end (see Figure 1).

### Applying tDCS

The total applied tDCS current was 4mA in the SE and ME montages; in the SE montage, all 4mA was applied through a single electrode while in the ME montage, 2mA was applied to the posterior STG/MTG junction and 1mA each was applied to the SMG and posterior IFG electrodes. The stimulation was applied using the multichannel MR compatible Neuroconn DC-MC device (Neurocare, Germany). The Neuroconn stimulator device was placed in the MR control room where the stimulation signals were generated. Three channels were programmed to concurrently generate 1mA and 2mA currents for the multi-electrode condition or 4mA in a single channel output for the single electrode condition (Figure 1a). In each of the concurrent tDCS-fMRI sessions, we alternated between the stimulation OFF and ON conditions, starting with the OFF condition. Three 4-minute epochs of stimulation were applied during the 24-minute resting state fMRI. Figure 1b shows the timing diagram of the stimulation epochs during one rs-fMRI acquisition of 24 minutes.

### Safety and tolerability

After each session of tDCS, we recorded safety and tolerability information. The skin and scalp location under the electrodes were inspected for any skin burns or other lesions. At the end of each session, we asked volunteers to indicate their tolerance of the noninvasive brain stimulation on a visual analog scale (VAS) with 0 and 10 as the endpoints where zero indicated that subjects tolerated the stimulation well and had no unusual sensations and 10 indicated that the stimulation session caused strong sensory experiences and was judged to be barely tolerable.

### Dynamic functional connectivity matrices

Although we recorded rs-fMRI with three stimulation epochs interspersed by non-stimulation epochs, in the current report we focus on the first stimulation epoch, especially the correlation matrices generated from 4D rs-fMRI image sets acquired immediately after the first stimulation onset. ML models designed in this case rely on feature vectors to perform classification. Hence, we used a vector representation of a dynamic functional connectivity (DFC) matrix as input feature vector. DFC is a method of calculating variations in functional connectivity between iterative samples of two regions generating a DFC matrix for a selected window length, then moving the time window by a step size to calculate the next DFC matrix which can also be termed as a sliding window functional connectivity. A DFC matrix for a selected time window is calculated in two steps-1) extract (or derive) the raw fMRI signal from the time-course for a region/s of interest (ROI) and 2) correlate that time-course signal from each ROI with all of the other ROIs of interest in a correlation matrix. We selected 1 minute as a window period and the step size of one acquisition volume (approximately 3 seconds). Figure 3 shows the data preprocessing pipeline including DFC calculation, window size and step size. Further, details on preprocessing parameters are provided in supplementary document (Table S1)

**Figure 3:**
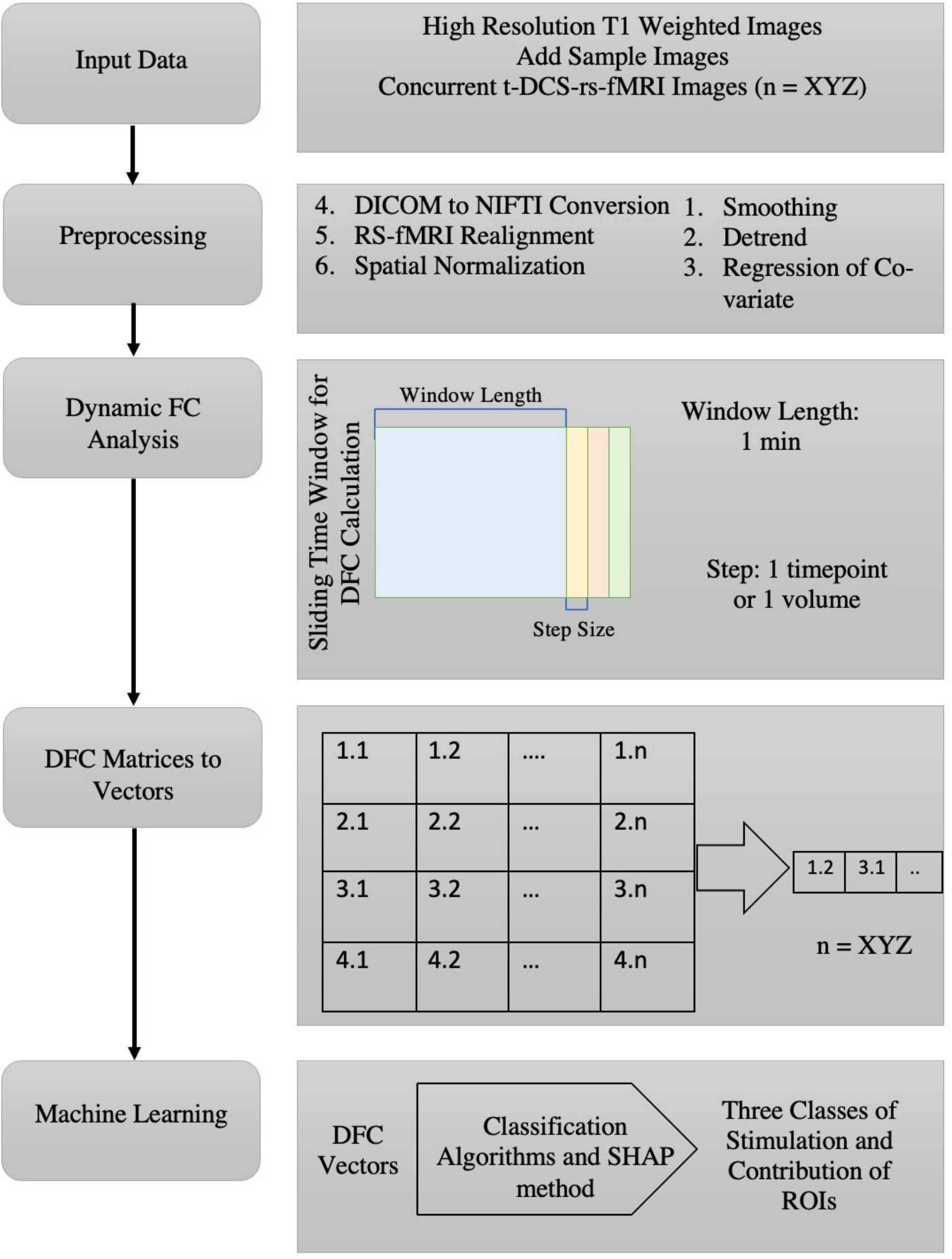
Data preprocessing flow-chart

A DFC for the no-stimulation conditions was calculated assuming the onset time of a “pretend-stimulation” as 3 minutes after the start of the acquisition to mimic the onset times of the SE and ME conditions. Data from the first 2 minutes of acquisitions after the real or pretend onset time was used to generate the DFC matrices and consequently the NS featured class vectors. We performed three sub-analyses to identify the shortest fMRI duration that can be used to classify the three different tDCS conditions. In each sub-analysis, a different number of DFC matrices were used as feature vectors. The number of DFC matrices used was either 5, 10, or 20, representing the first 15, 30, or 60 seconds of fMRI data. These DFC matrices were used to generate the featured class vectors which were then grouped into two categories: single electrode stimulation onset, and multi-electrode stimulation onset. The groups of data from the stimulation sessions were then individually grouped with the NS featured class vectors resulting in a cumulative dataset of three classes. Although the stimulation was applied for 4 minutes which resulted in 80 DFC matrices, only first minute of the rs-fMRI data was used to perform classification.

### Selection of ROI pairs

The standard Harvard-Oxford atlas [35–38]with 112 regions of interest was used to calculate a 112 × 112 DFC correlation matrix. Because of the structure of the correlation matrix, each value is replicated and the values on the diagonal axis are all ‘1.’ We converted each DFC matrix into a vector of 1 × 6216 unique ROI pairs.

The large size of the feature vector can increase time requirements for preprocessing and classification. To develop a real time feedback system and reduce the data processing time, we devised a strategy to reduce the number of ROIs based on neuroanatomy and focus on the cortical regions associated with AF-network. ROIs were drawn on MNI-space T1-weighted images and a total of 38 equally sized spheres (16 mm in diameter) intended to sample previously-identified cortical nodes of the AFN as well as the cortical nodes of the Inferior Longitudinal Fasciculus (ILF) network, which was chosen as a control network in both hemispheres. This resulted in a 38 × 38 DFC correlation matrix with a feature vector size of 1 × 703. To reduce the size of the feature vector even further, we reduced the number based on their importance and closeness to the nodal cortical endpoints of the AF-network to generate three groups with a total of 26, 22, 16 ROIs corresponding to the vector sizes 1 × 325, 1 × 231, and 1 × 120. A list of these custom ROIs and ROI groups is provided in the supplementary document (Table S2).

### fMRI to feature vector calculation

We used a graph theoretical network analysis toolbox called GRETNA for preprocessing and calculation of DFC matrices [40]. First, DICOM images were converted to 3D (T1w images) and 4D (rs-fMRI) NIfTI (Neuroimaging Informatics Technology Initiative) datasets using dcm2niix [41]. Since the BOLD signal shows some T1-saturation effects in the first volume acquisitions, we excluded the first minute of the rs-fMRI data from the analysis. This was followed by preprocessing steps such as slice timing correction, realignment of rs-fMRI images, spatial normalization to the standard MNI space using the dartel segmentation method, smoothing, detrending fMRI data, and regressing out covariates such as white matter signal, CSF signal and head motion. The regression was carried out using a published strategy [42]. After the DFC matrices were generated, we used a custom program developed in MATLAB to get the feature vector for each participant and each set of ROIs to be used in the ML part.

### Machine learning approach

#### Dataset preparation

The prepared dataset for this study consists of data from 33 participants who completed a total of 69 sessions which included 20 no-stimulation sessions, 25 single-electrode sessions, and 24 multi-electrode sessions. In order to ensure that the classification models used in this study were able to train effectively and were tested on an unseen data to assess performance of the trained model, the dataset was divided into a training set (80% of the data) and a testing set (20% of the data). Additionally, we ensured that data from a particular participant was not split between testing and training data which lead to 26 datasets used for training and 7 datasets (not seen by the model) used for testing. However, due to the fewer number of sessions included in the dataset, there was a risk that the classification models would be overfit to the training data and not generalize well while performing on the testing data [43]. This means that the generalized learning of mapping from inputs to outputs is avoided, and the models could learn the specific inputs and the associated output results [44]. To mitigate this risk, we took two approaches: up-sampling the training data and adding noise to the training data [45]. Up-sampling involved an increase in the number of training examples [46,47], while adding noise across the training data involved randomly swapping the values of multiple features for a subset of the data points. These measures were taken to make the classification models more robust and better able to generalize their predictions regarding new data.

#### Classification algorithm

The machine learning (ML) models random forest, k neighbors’ classifier, naive bayes, decision tree classifier, gradient boosting classifier were used on the feature vectors from the DFC matrices. The objective of the ML models is to classify the DFC matrices into three categories: NS, SE, and ME [48].

Ten-fold cross-validation was deployed while training the models. The use of multiple models has helped us compare performance across different models along with a varying number of volumes.

#### Independent validation

To validate the results of the classifier and to assess the performance of the classification model, we focused on three parameters: Receiver Operating Characteristic (ROC) score, Matthews Correlation Coefficient (MCC), and Accuracy (based upon the test dataset). The ROC score shows the relationship between sensitivity and specificity for every possible cut-off when performing a combination of predictions. The MCC score calculates the difference between the actual values and predicted values of the classification which is equivalent to the chi-square statistics for a 2 × 2 matrix [49]. While evaluating the models using different parameters, we have also considered each model’s sensitivity and specificity. Accuracy was calculated using the ratio of true positives and true negatives against all predictions.

#### Model interpretation

There are various methods to estimate the importance of the attributes. In tree-based models like RF and decision tree, attribute importance is pre-assigned which greatly influences the output. The contribution of an attribute is completely different from its importance towards the output. The importance is based on the quantitative approach that determines which attributes significantly drive the model’s performance. Whereas the contribution of the attributes helps to understand an intuitive explanation for the output, beyond just identifying what attributes strongly affect the classification performance.

In this study, we have used the Gini impurity decrease to interpret RF model. We can determine how much a particular attribute contributes toward the average decrease in the error of the model classification. Each value from the feature vector represents a correlation between ROI pairs and Gini impurity decrease and helps us identify the importance of each ROI pair or an attribute. However, this method is insufficient in explaining how individual attributes affect the prediction.

Therefore, we also used the SHAP method to determine quantitatively how each attribute contributes to the RF model’s performance [50,51]. The SHAP uses the game theory concept to calculate the contribution of each of the attributes combined with the prediction model and explanation model using the various methods. Using the SHAP method we can determine the contribution of each attribute to the model classification.

#### Proof-of-concept real-time feedback system

The various classification models described above will be tested to identify the ML models’ capable of identifying the effects of ME and SE stimulation on the fMRI data. Additionally, SHAP method and Gini impurity decrease interpretation will be used to identify the top ROI pairs that contribute strongly towards this classification.

This was done as follows: whenever a new fMRI dataset was presented to the system, it was processed to calculate the DFC matrices using GRETNA software which would then be converted to the feature vectors and fed as input to the classification model. The pre-trained classification model would inform if the fMRI data showed responses that correspond to the effects generated with ME and SE stimulation. Additionally, top contributing ROI pairs can be compared against top contributing pairs generated while training the model. Upon receiving new data, the model first generates predictions based on its previous training. Then, the dataset is expanded by incorporating the new data and human-verified results from the model’s predictions. This updated dataset is used to retrain the model, ensuring that it is able to continuously improve its learning.

This proof-of-concept feedback system is designed to help in the evaluation of unknown (newly acquired) fMRI data using classification model and corresponding interpretation indices which can inform about the stimulation and similarity of the top contributing ROI pairs with the ROI pairs that strongly influence the classification in validation. Using this information, stimulation parameters can be adjusted to invoke the desired responses in fMRI.

## Results

### Evaluation of classification performance

Classification performance was evaluated using the feature vectors generated with the first 15, 30, and 60 seconds of the rs-fMRI data immediately after stimulation onset. Best classification performance was observed with the first minute of data which are detailed below. Results with the first 15 seconds and first 30 seconds of rs-fMRI data are presented in the supplementary document.

### ROC and MCC score

The ROC score was generated using the true positive rates and false positive rates of the models. The MCC score is used as a measure of the quality of binary classifications and has been generated using the formula based on true positives, false positives, true negatives, and false negatives. In table 1, we represent the ROC and MCC scores for each of the models. A score of 1.0 for the ROC indicates the model performs ideally. In the case of the k-nearest neighbor (knn) classifier, the ROC was consistently 0.90 (except when 112 ROIs or 16 ROIs were used) or higher when compared to the other models, this displayed the best performance. Although the Random Forest Classifier came close to the values of the ROC curve it was consistently lower compared to knn. A score of 1.0 for the MCC indicates the perfect agreement between the prediction and observation. MCC values for the knn classifier were close to 0.85 when 38, 26 and 22 ROIs were used to calculate feature vectors. Similar to ROC results, MOC values with random forest classifier were close to the knn performance but were slightly lower. Logistic regression, decision tree, and naïve Bayes classifiers’ performance was inferior to the knn or random forest classifiers irrespective of the number of ROIs used to generate feature vectors.

**Table 1:** ROC and MCC across different ROIs for the classification shown by various models

| Model | No. of ROIs - 112 |  | No. of ROIs - 38 |  | No. of ROIs - 26 |  | No. of ROIs - 22 |  | No. of ROIs - 16 |  |
| --- | --- | --- | --- | --- | --- | --- | --- | --- | --- | --- |
|  | ROC | MCC | ROC | MCC | ROC | MCC | ROC | MCC | ROC | MCC |
| K Neighbors Classifier | 0.6822 | 0.6402 | **0.9055** | **0.8593** | **0.9099** | **0.8697** | **0.9247** | **0.8459** | **0.8898** | **0.8213** |
| Random Forest Classifier | **0.7321** | **0.6789** | 0.8803 | 0.8207 | 0.8889 | 0.8124 | 0.9012 | 0.8411 | 0.8646 | 0.7984 |
| Logistic Regression | 0.7091 | 0.6211 | 0.8519 | 0.7794 | 0.8697 | 0.7985 | 0.8899 | 0.7456 | 0.7836 | 0.6732 |
| Decision Tree Classifier | 0.633 | 0.5154 | 0.7553 | 0.6338 | 0.7375 | 0.6741 | 0.8838 | 0.7982 | 0.7533 | 0.6173 |
| Naive Bayes | 0.654 | 0.4879 | 0.72 | 0.5856 | 0.7356 | 0.6569 | 0.7099 | 0.6428 | 0.5596 | 0.3215 |

### Classification accuracy

In table 2, we show the overall classification accuracy of different models with varying sets of ROIs. It is observed that the overall accuracy improved when the number of ROIs were reduced from 112 to lower numbers, and it was highest [0.9247] when using the dataset generated using 22 ROIs and k-nearest neighbor (knn) classifier.

**Table 2:** Accuracy across different ROIs for the classification shown by various models

| Model | No. of ROIs - 112 | No. of ROIs - 38 | No. of ROIs - 26 | No. of ROIs - 22 | No. of ROIs - 16 |
| --- | --- | --- | --- | --- | --- |
| K Neighbors Classifier | 0.683 | **0.9056** | **0.9099** | **0.9247** | **0.8833** |
| Random Forest Classifier | **0.731** | 0.88 | 0.8889 | 0.9012 | 0.8689 |
| Logistic Regression | 0.71 | 0.8518 | 0.8697 | 0.8899 | 0.7869 |
| Decision Tree Classifier | 0.63 | 0.7556 | 0.7375 | 0.8838 | 0.75 |
| Naive Bayes | 0.6544 | 0.722 | 0.7356 | 0.7099 | 0.5574 |

Figure 4 shows the ability of top 5 performing models to classify the three stimulation conditions with feature vectors extracted from the rs-fMRI data with 22 ROIs. For each model, bar graphs represent classification accuracy for each class. It can be observed that in some cases, the knn classifier is not the one which classifies with highest accuracy. However, it is our best performing model because when taken overall, it shows highest accuracy. Figure 4 also shows that the random forest model classifies the NS and SE with the highest accuracy values while the knn model classifies the ME category with the highest accuracy. Additionally, figure 5 shows confusion matrices for knn and random forest models generated for the dataset with 22 ROIs and 26 ROIs.

**Figure 4:**
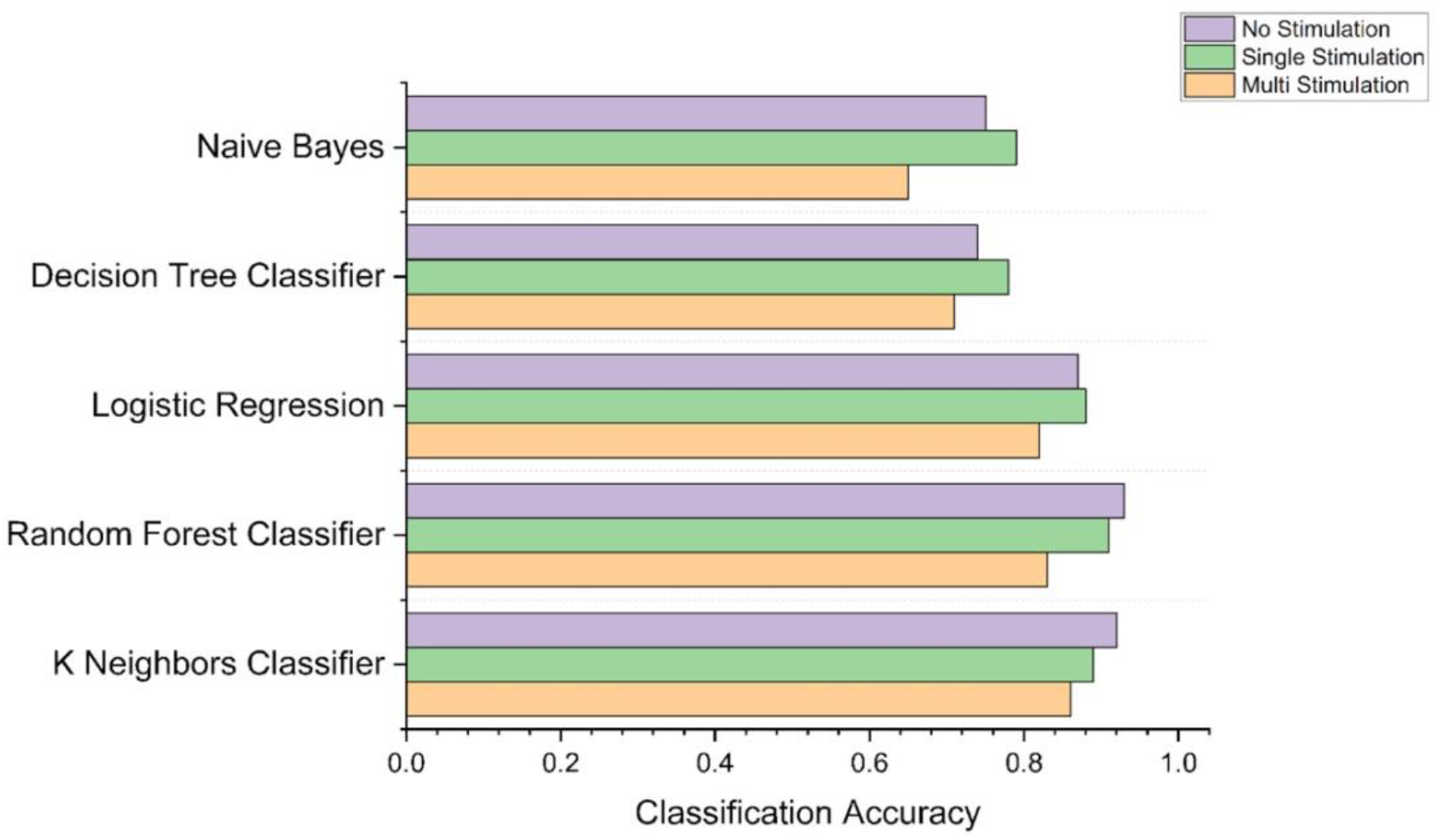
Accuracy of models to classify different stimulations

**Figure 5:**
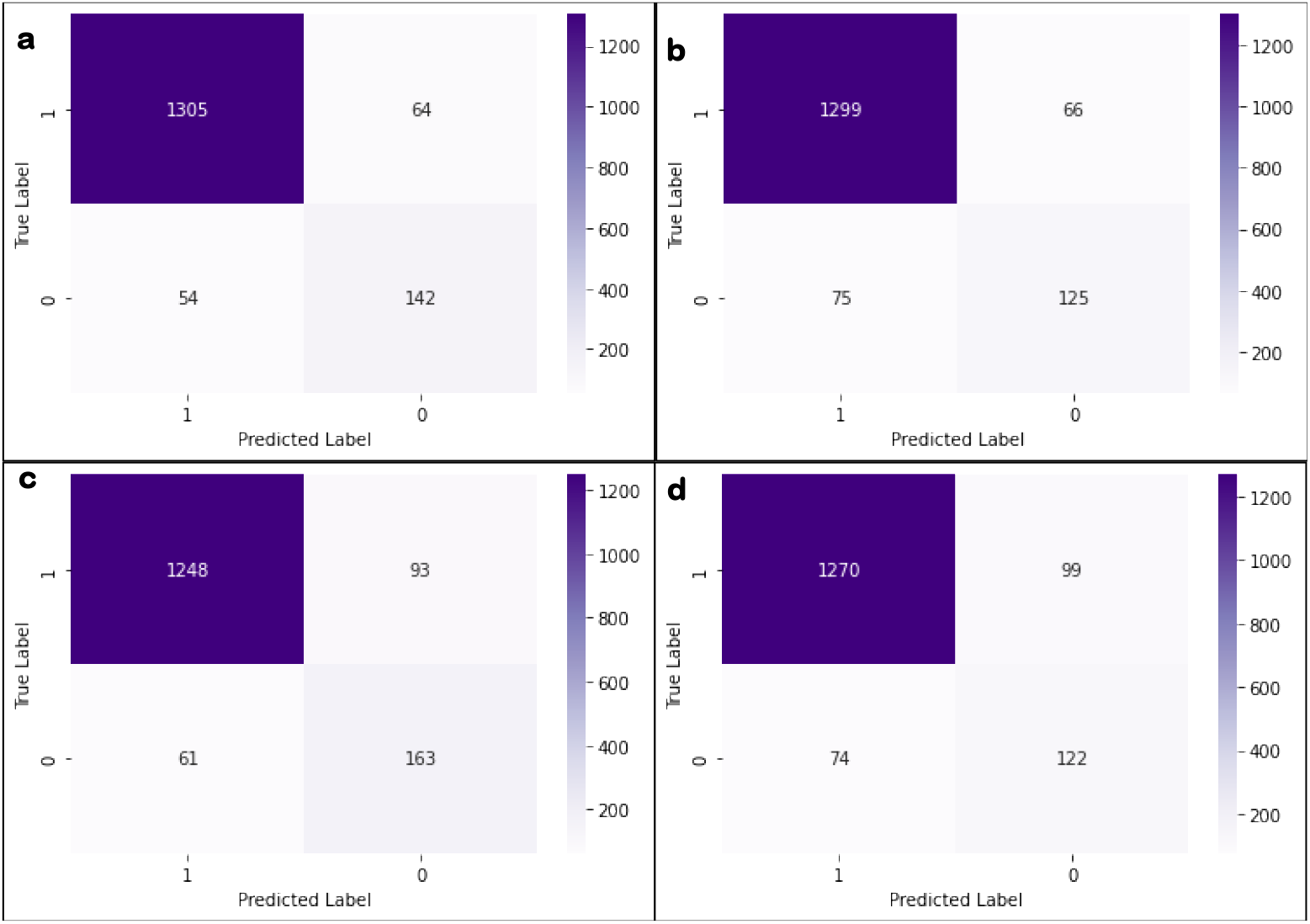
Confusion matrices for the prediction of knn model on the 22ROIs (a) and 26 ROIs(b) dataset. Confusion matrices for the prediction of random forest model on 22 ROIs (c) and 26 ROIs (d) dataset

### Analysis of the contributions and importance of attributes

For further analyses, we have selected the 22 ROI and 26 ROI sets because these sets showed the best performance in machine learning predictions and took comparatively less computational time. Since Random forest algorithm was the best performing decision tree model, we evaluated contribution and importance of attributes using Gini impurity decrease and SHAP. The Gini impurity decrease can be used to evaluate the purity of the nodes in the decision tree, while SHAP can be used to understand the contribution of each feature to the final prediction made by the model. This helped in identifying any biases or patterns in the model’s predictions and understand why the model is making certain decisions.

In figure 6, the importance plot shows the RF model’s ability to classify based on the Gini impurity decrease for each attribute using 22 ROIs and 26 ROIs, respectively. The top ROI pair from the data with 26 ROIs has the Gini impurity decrease of 0.174, and subsequently, the tenth most important pair has the Gini impurity decrease of 0.031. The top ROI pair from the data with 22 ROIs has the Gini impurity decrease of 0.246, and subsequently, the tenth most important pair has the Gini impurity decrease of 0.019. Although the sum of the Gini impurity decrease for all pairs is equal to 1, the top 5 ROI pairs in the 26 ROIs and 22 ROIs contribute more than 50% towards it.

**Figure 6:**
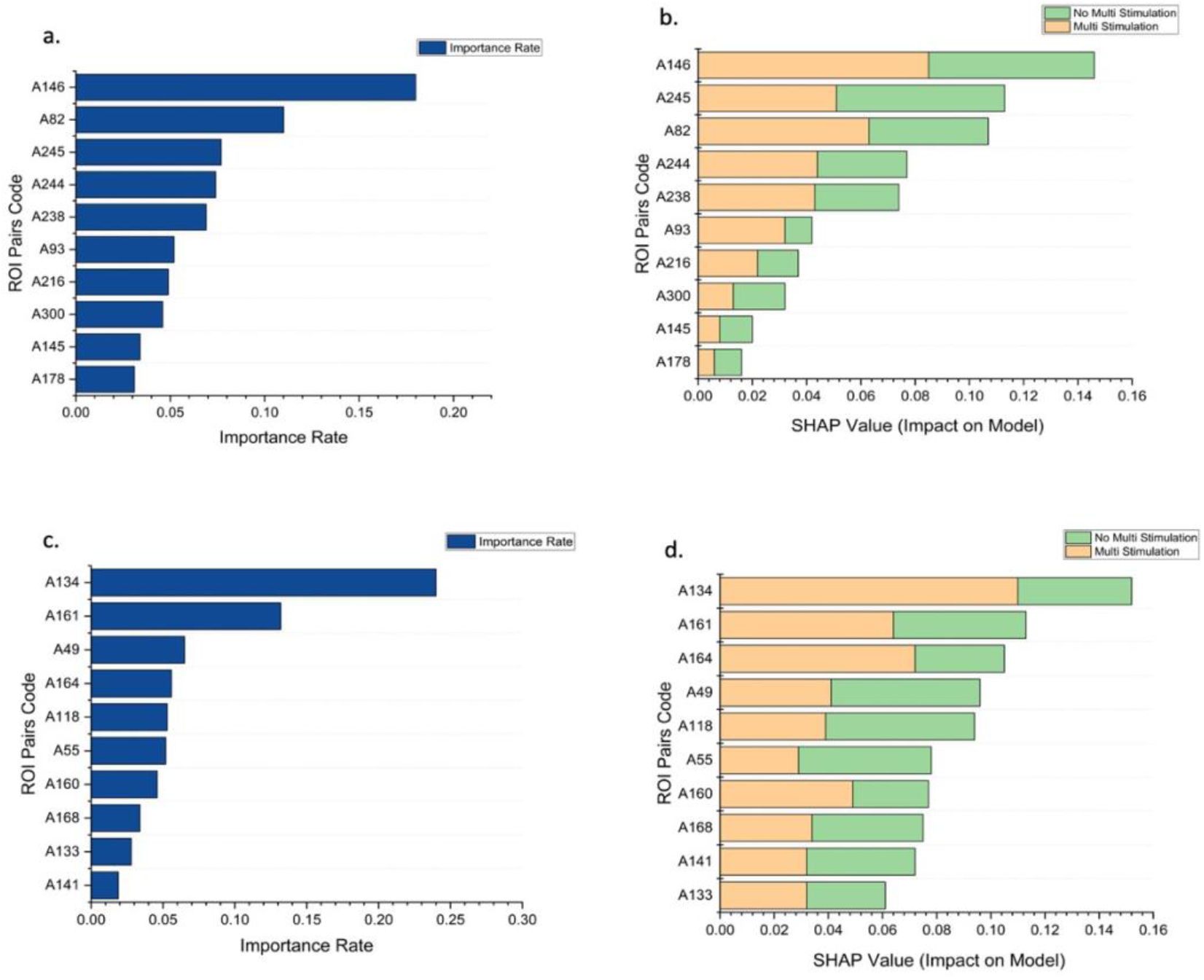
Importance rate and contribution of ROI pairs towards classifying ME stimulation using Gini impurity decrease and SHAP value: (a & b) 26 ROIs and (c & d) 22 ROIs. Table S7 and S8 in supplementary document lists the ROI pairs associated with the codes.

The importance rate doesn’t tell us about the contribution of each attribute towards a specific output. The SHAP summary plot’s introduction helps in performing an in-depth analysis. From the figure 6b, it is observed that the top pair from the data with 26 ROIs has the highest contribution of 0.086 towards the model’s output of classifying a vector into ME. In contrast, the second ROI pair has the highest contribution of 0.081 towards the model’s output of classifying a vector into not ME. Similarly, figure 6d represents the SHAP values for pairs of 22 ROIs. It is observed that the top pair has the highest contribution of 0.118 towards the model’s output of classifying a vector into ME. In contrast, the fourth and fifth ROI pairs have the highest contribution of 0.055 towards the model’s output of classifying a vector into not ME. This shows how each ROI pair can affect a specific output by the RF model.

### Time Taken for computation for processing of ROIs

We used a computer with 64 gigabytes of RAM, NVIDIA GeForce RTX 2080 of 4 gigabytes, and AMD Ryzen 9 5900X CPU with UBUNTU 18.04 operating system. The GRETNA program was installed on MATLAB 2021b (MathWorks, Natick, MA) which was used to carry out the entire processing pipeline including calculation of the DFC matrices. Time required for preprocessing steps was 11 minutes and 26 seconds and it did not differ with number of ROIs. In contrast, the calculation of DFCs showed exponential decrease in time required with a decrease in the number of ROIs. In figure 7, we see that the time taken for the computation of DFC calculation for different numbers of ROIs has inverse exponential relation with a decrease in the number of ROIs. The maximum time taken to calculate the DFC is 6minutes and 18 seconds with 112 ROIs whereas it came down to just 27 seconds when calculated for 16 ROIs.

**Figure 7:**
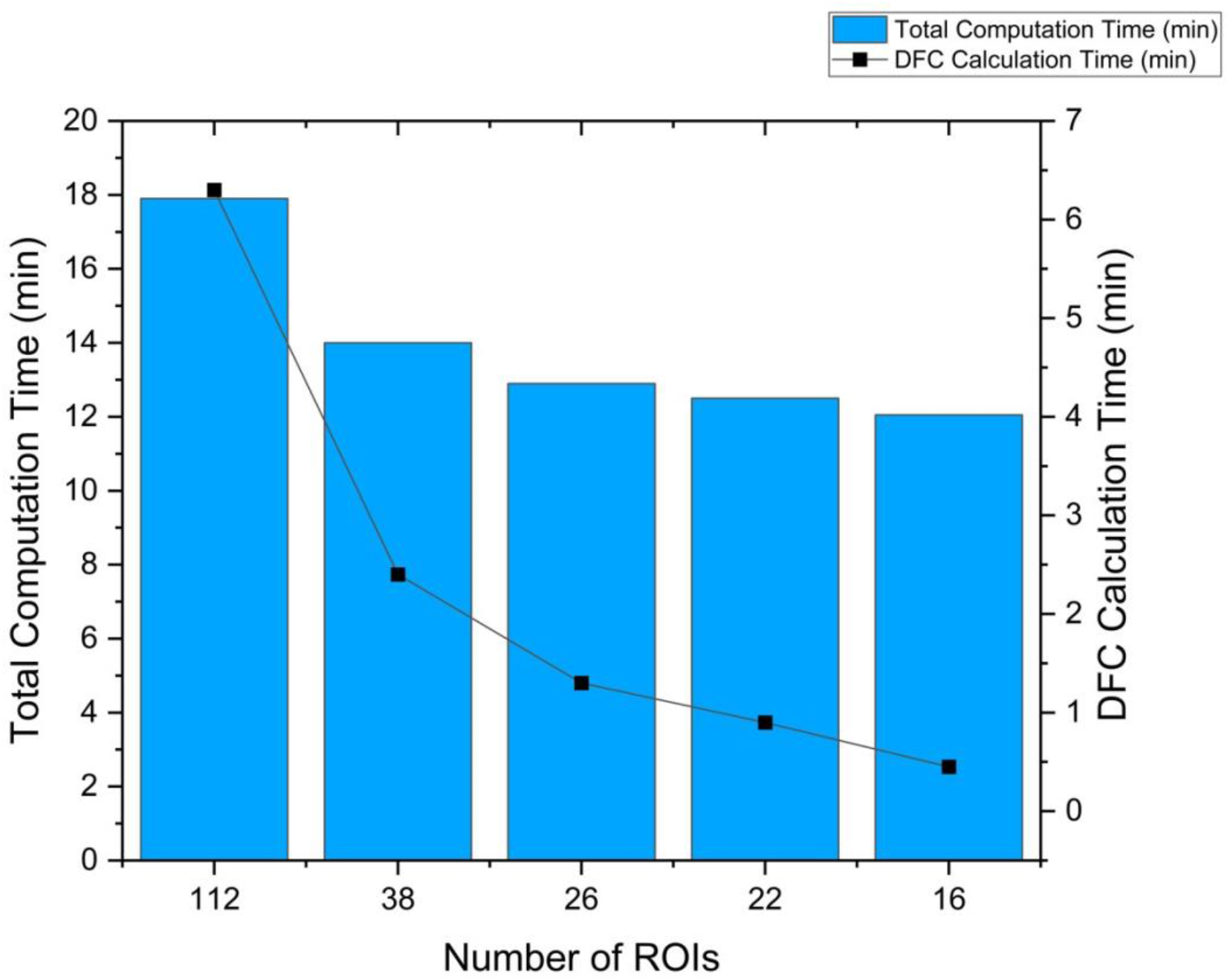
Time taken for data processing up to prediction, varying the number of ROIs used

### Evaluating the accuracy of real-time feedback system

In this study, we applied two models - best performing pre-trained model (knn) and best performing tree-based model (random forest) to classify the test dataset into various conditions. We computed the Gini impurity decrease and SHAP methods over the RF model to identify the contribution and importance of each ROI pair.

In each test case, we found that the similarity of the top 10 ROI pairs is consistently higher than 60% with the averaged classified list of ROI pairs for ME class. In each independent dataset’s predictions, we identified that the top 2 contributing ROI pairs were SMA1_R – iPreCG_R and ParOp_R – IFG3_R; those two pairs match the corresponding averaged classified list of pairs, as shown in figure 8.

**Figure 8:**
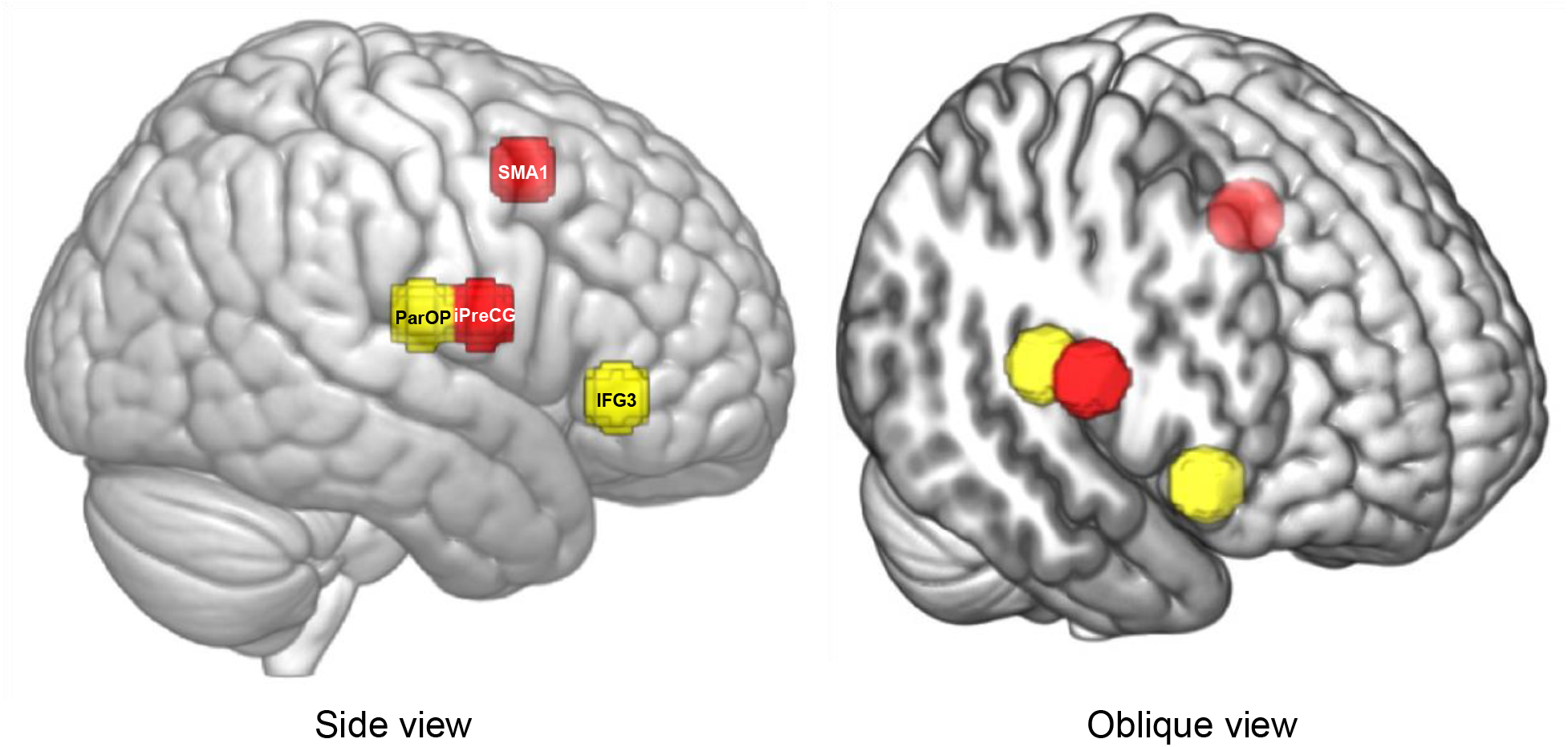
The top 2 ROI pairs contributing towards the classification of multi-electrode stimulation

The total time taken in this process of prediction of one test dataset was close to 18 minutes (12 minutes of preprocessing and 6 minutes for feature extraction and classification). Such a short time would allow researchers to continue the experiment with a 30 min break to determine engagement of a targeted brain region and adjust the electrodes if necessary to achieve true engagement.

### Evaluation of Safety and tolerability

The study using the high dose tDCS stimulation caused no significant adverse effects among the participants. None of the participants experienced severe headaches, seizures, neurological impairments, skin burns, or any hospitalizations that were directly related to the stimulation.

In addition, stimulation was well tolerated by all the participants. None of the participants stopped the experiment due to the stimulation being intolerable (=VAS scores 9 and 10). Figure 9 shows the tolerability scores for SE and ME sessions, and a T-test between the scores showed no significant difference (p = 0.8455).

**Figure 9:**
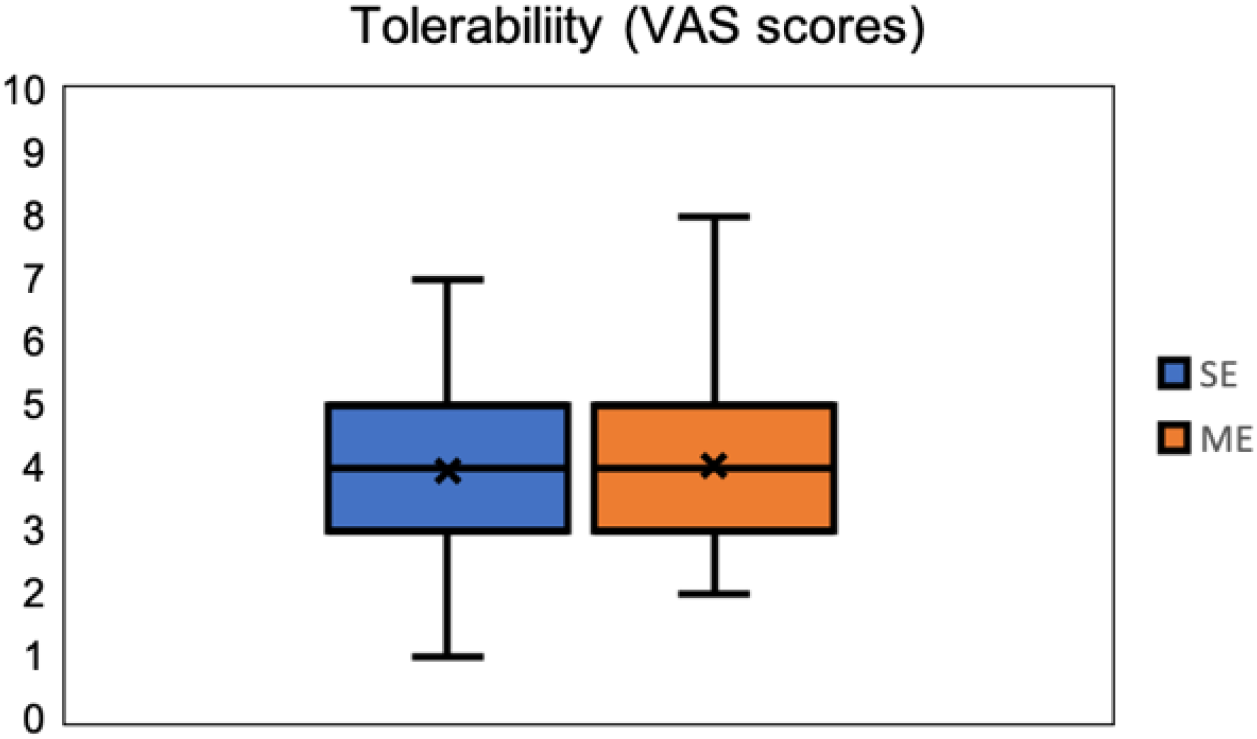
VAS tolerability scores (min=0, max=10) of all sessions of all participants

## Discussion

We developed a classification method that used a complex 4-D fMRI data set acquired during concurrent tDCS-fMRI experiments to identify whether or not (1) stimulation was applied and (2) electrode montage had an effect on network engagement. A multi-electrode montage targeting nodal points of access more reliably engaged the AFN than a single electrode montage or a no-stimulation condition. Additionally, we developed a proof-of-concept feedback system using machine learning methods to determine quickly and efficiently whether or not the best montage modulated the targeted brain network or not. We presented a unique and innovative approach to use machine learning applied to 4-D concurrent tDCS-fMRI data to classify tDCS effects based on stimulation montage and to generate a real-time feedback system to verify engagement of the targeted network.

Advancement of the novel machine learning approaches, increase in computational processing speed, and data analysis techniques have fueled the use of artificial intelligence (AI) techniques to solve multi-dimensional data-intensive problems [52]. The use of AI has shown promising results in different healthcare applications including X-ray & CT evaluation, development of antibiotic drugs, cancer classification, biomarker detection, and diagnosis [53]. Only a limited number of research studies have evaluated the use of AI to understand the effects of targeted tDCS parameters and their topical specificity. Al-Kaysi and colleagues used baseline EEG spectra and mood-cognition evaluation measures from 10 participants’ data to predict if tDCS will positively modulate the mood [30]. Kajimura and colleagues showed that functional MR images acquired after active stimulation differed compared to sham; however, this study suffered from two limitations: there were few participants in each group (N=12) and applying tDCS outside the scanner which means that the study investigated effect of tDCS in different physical conditions [31]. In other studies, ML methods were used to perform binary classification in complex data, outcome variables and their relationship to the stimulation, and how other baseline factors affect the outcome variables [30,32–34]. All of these studies, used ML methods to either understand/investigate effects of the stimulation in two groups namely ‘Stimulation’ and ‘No-Stimulation.’ Our study was more complex than previous studies in two aspects, first, we used concurrent tDCS-fMRI allowing us to record real time effects of stimulation and we applied ML methods to perform classification of three categories including two different stimulation montages. The classification accuracies demonstrate the ability of the ML model to detect if stimulation was applied or not. Additionally, the ML model reliably classifies condition into ME, SE, and NS stimulation categories with the ME category achieving the highest accuracy.

Our secondary goal was to develop a real-time feedback system that uses the shortest fMRI acquisition time to determine if the targeted network has been engaged. Real-time has to be defined for each system as it is a subjective measure, in this case we defined it as half an hour or less. This period of time might be reasonable to determine whether or not the experiment can be continued, or electrodes have to be repositioned. The alternative approach without such a fast feedback system would be the collection of an entire dataset, the off-line analysis of such a dataset to determine if a network has been engaged or not, and a decision whether or not the experiment that relied on a true network engagement would have to be repeated again. A new electrode montage would lead to a new experiment several days later and the same cycle would start over again. Our proposed feedback system of reducing the sets of ROIs for computing efficiency and accuracy, although potentially increasing the initial fMRI session’s duration, would reduce the need for subsequent sessions and time and effort to analyze datasets that might not show an engagement of the targeted system. As shown in the results section, ROI sets with 22 ROIs perform classification at overall accuracy close to 90% and time required for feature calculation close to 1 minute compared to 6 minutes required when using 112 ROI annotated over the whole brain with total computation time close to 18 minutes. In addition to identifying how many ROIs would be optimal, we also investigated which classification model would provide the best possible accuracy, ROC, and MOC scores. Results suggest that KNN model performs at the highest accuracy of 92% closely followed by the random forest model with the accuracy of 90%.

The proof-of-concept feedback system used DFC between ROI pairs as an individual feature; hundreds of such features were used as input for the ML method. However, identifying the anatomically relevant ROIs not only improved the accuracy, reduced time required, but also improved the possibility of using ML method as a feedback system by allowing us to identify the top contributing features using the Gini impurity decrease and the SHAP method. The top 5 out of 231 features contributed close to 50% towards the classification.

Testing of the independent fMRI dataset with the feedback system showed that the top DFC of the two ROI pairs contributing towards the classification into stimulation type highly matches with the top two contributing features generated from the group analysis. The top two ROI pairs that are identified as strong contributors to the ME classification were ‘SMA1_R – iPreCG_R’ and ‘ParOp_R – IFG3_R.’ (see Figure 8 for details of the anatomical location of these spherical ROIs). These ROI pairs represent nodal regions and relay regions of the right AFN, connected via the AF tract. Although the ROIs were anatomically defined and chosen to be of equal size, they were specifically determined by the investigators to examine the AFN and ILF. In the future more generic ROIs derived either from atlases or defined using resting-state discrete atoms [54] or larger established resting-state networks could be tested. The ME montage is designed to stimulate multiple nodal regions of the right AFN with the goal to increase connectivity between them as well as other related network regions. The results of our classification analysis suggest that it is the change in connectivity between regions belonging to this network that enhances the classification output. Further, 60% of the top 10 features that contribute towards the classification are common across participants. The number 60% is not just more than a random chance, but it is a significantly stronger measure of commonalities in the response across participants as these pairs would contribute more than 50% towards classification of the participant to a particular stimulation type.

There are certain limitations where the ML approach can be improved. The first one being the time required for preprocessing, we are exploring if improving computing power and using high speed memory devices will reduce the time required, additionally, we are also exploring the possibility of performing these operations over high performing computing machines to harness their processing speed. This would allow us to shorten the period from concurrent tDCS-fMRI acquisition to the feedback on targeted engagement to the experimenter. Second, the current proof-of-concept feedback system could be further optimized by the use of an MR-compatible multiplexed electrode system that can change where the current is applied to achieve network effects using a pre-mounted array of electrodes making it less likely that a participant would have to be moved out of and back into the scanner again and adjusting location of the electrodes.

In conclusion, machine learning was developed to identify the best electrode montage to show the most engagement of a targeted brain network spanning multiple lobes of the brain with high accuracy. ROI pairs whose DFC values contributed strongly towards the multi-class classification were identified in the order of importance. The top 5 ROI pairs, out of 231 ROI pairs, have a cumulative importance of 50% and are all connected to the AFN. Furthermore, a prototype of a machine learning real-time feedback system was introduced which would only require 1 minute to extract the DFC calculation-parameters and 1 minute for ML calculations.

## Acknowledgement

This research was supported by an NIH BrainInitiative grant (<u>RO1MH111874</u>). Dr. Schlaug also acknowledges support from U01NS102353. AS acknowledges support from UMass Initiative on Neurosciences Inspiration Awards for Neuroscience and Technology. SM acknowledges the support from IALS summer internship program. We are thankful for the generous support that Klaus Schellhorn from neuroConn has provided to us with making a state-ofthe-art multi-channel MR DC stimulator available to us and for being available for any trouble shooting over the years. We would also like to thank Elena Bliss (MR technologists UMass Amherst) for her support with the concurrent tDCS-MR acquisitions.

