## SupplementaryInformation for "Identifying the engagement of a brain network during a targeted tDCS-fMRI experiment using a machine learning approach"

### Supplementary Information

Table S1 presents data on the specific settings or options that were chosen in the process of calculating dynamic functional connectivity (DFC) for each session using the GRETNA software. The GRETNA software is a tool that allows researchers to analyze functional magnetic resonance imaging (fMRI) data and calculate DFC.

Table – S1: Information regarding specific settings used in the GRETNA software while extracting the DFCs.

| **Parameters** | **Specific Input** |
|---|---|
| Slice Order | Alternating in the plus direction (starting at odd) |
| Reference Slice | Middle Slice |
| Normalization Strategy | DARTEL |
| Voxel Sizes (mm) | [1 1 1] |
| FWHM Size (mm) | [4 4 4] |
| Detrending Order | Linear and Quadratic |
| Regress Out Covariates | White Matter Signal, CSF Signal, Head Motion Strategy: Friston – 24 Parameters |
| Interpolation Strategy | Nearest Interpolation |
| Fisher’s Z Transformation | True |
| Sliding Window Step | 1 |
| Sliding Window Length | Varies according to TR |

Table S2 represents the ROIs which were identified to evaluate the effects of tDCS.

Table – S2: ROI sets used for analysis and prediction of DFCs of tDCS.

| No. | 16 ROIs | 22 ROIs | 26 ROIs | 38 ROIs |
|---|---|---|---|---|
| 1 | IFG1_L | IFG1_L | IFG1_L | IFG1_L |
| 2 | IFG1_R | IFG1_R | IFG1_R | IFG1_R |
| 3 | IFG3_L | IFG3_L | IFG3_L | IFG2_L |
| 4 | IFG3_R | IFG3_R | IFG3_R | IFG2_R |
| 5 | IFG4_L | IFG4_L | IFG4_L | IFG3_L |
| 6 | IFG4_R | IFG4_R | IFG4_R | IFG3_R |
| 7 | IFG5_L | IFG5_L | IFG5_L | IFG4_L |
| 8 | IFG5_R | IFG5_R | IFG5_R | IFG4_R |
| 9 | occMed_L | occMed_L | iPreCG_L | IFG5_L |
| 10 | occMed_R | occMed_R | iPreCG_R | IFG5_R |
| 11 | SMG1_L | ParOper_L | occMed_L | iPreCG_L |
| 12 | SMG1_R | ParOper_R | occMed_R | iPreCG_R |
| 13 | SMG3_L | SMA1_L | ParOper_L | mPreCG_R |
| 14 | SMG3_R | SMA1_R | ParOper_R | mPreCG_L |
| 15 | tempMed_L | SMA2_L | pSTG_L | occLat_L |
| 16 | tempMed_R | SMA2_R | pSTG_R | occLat_R |
| 17 | | SMG1_L | SMA1_L | occMed_L |
| 18 | | SMG1_R | SMA1_R | occMed_R |
| 19 | | SMG3_L | SMA2_L | ParOper_L |
| 20 | | SMG3_R | SMA2_R | ParOper_R |
| 21 | | tempMed_L | SMG1_L | pMTG_L |
| 22 | | tempMed_R | SMG1_R | pMTG_R |
| 23 | | | SMG3_L | pSTG_L |
| 24 | | | SMG3_R | pSTG_R |
| 25 | | | tempMed_L | SMA1_L |
| 26 | | | tempMed_R | SMA1_R |
| 27 | | | | SMA2_L |
| 28 | | | | SMA2_R |
| 29 | | | | SMG1_L |
| 30 | | | | SMG1_R |
| 31 | | | | SMG_L |
| 32 | | | | SMG2_R |
| 33 | | | | SMG3_L |
| 34 | | | | SMG3_R |
| 35 | | | | tempLat_L |
| 36 | | | | tempLat_R |
| 37 | | | | tempMed_L |
| 38 | | | | tempMed_R |

Tables S3+S4 represent the sub-analyses: data from only the first 5 DFC matrices (=15 seconds of scan time) is included.

Table – S3: ROC and MCC across different ROIs for the classification shown by various models

| Model | No. of ROIs - 112 | | No. of ROIs - 38 | | No. of ROIs - 26 | | No. of ROIs - 22 | | No. of ROIs - 16 | |
|---|---|---|---|---|---|---|---|---|---|---|
| | ROC | MCC | ROC | MCC | ROC | MCC | ROC | MCC | ROC | MCC |
| K Neighbors Classifier | **0.517** | **0.478** | 0.511 | 0.486 | **0.569** | **0.533** | 0.444 | 0.422 | **0.513** | 0.451 |
| Random Forest Classifier | 0.438 | 0.422 | 0.487 | 0.462 | 0.532 | 0.477 | 0.321 | 0.291 | 0.451 | 0.422 |
| Logistic Regression | 0.467 | 0.443 | 0.511 | 0.487 | 0.542 | 0.522 | 0.431 | 0.397 | 0.398 | 0.355 |
| Decision Tree Classifier | 0.458 | 0.421 | **0.531** | **0.498** | 0.544 | 0.531 | **0.578** | **0.552** | 0.512 | **0.483** |
| Naive Bayes | 0.358 | 0.347 | 0.421 | 0.396 | 0.317 | 0.289 | 0.346 | 0.331 | 0.341 | 0.321 |

Table – S4: Accuracy across different ROIs for the classification shown by various models

| Model | No. of ROIs - 112 | No. of ROIs - 38 | No. of ROIs - 26 | No. of ROIs - 22 | No. of ROIs - 16 |
|---|---|---|---|---|---|
| K Neighbors Classifier | **0.541** | **0.522** | **0.612** | 0.438 | **0.561** |
| Random Forest Classifier | 0.459 | 0.517 | 0.577 | 0.362 | 0.431 |
| Logistic Regression | 0.431 | 0.468 | 0.442 | 0.418 | 0.351 |
| Decision Tree Classifier | 0.437 | 0.521 | 0.573 | **0.592** | 0.485 |
| Naive Bayes | 0.328 | 0.385 | 0.428 | 0.366 | 0.342 |

Tables S5+S6 represent the sub-analyses when data included is from the first 10 DFC matrices (=30 seconds).

Table – S5: ROC and MCC across different ROIs for the classification shown by various models

| Model | No. of ROIs - 112 | | No. of ROIs - 38 | | No. of ROIs - 26 | | No. of ROIs - 22 | | No. of ROIs - 16 | |
|---|---|---|---|---|---|---|---|---|---|---|
| | ROC | MCC | ROC | MCC | ROC | MCC | ROC | MCC | ROC | MCC |
| K Neighbors Classifier | 0.415 | 0.387 | 0.411 | 0.375 | 0.525 | 0.475 | 0.483 | 0.468 | 0.397 | 0.365 |
| Random Forest Classifier | 0.441 | 0.435 | 0.475 | 0.461 | 0.379 | 0.388 | 0.477 | 0.451 | 0.308 | 0.287 |
| Logistic Regression | 0.539 | 0.527 | 0.531 | 0.487 | 0.418 | 0.387 | 0.511 | 0.495 | **0.629** | **0.572** |
| Decision Tree Classifier | 0.431 | 0.429 | 0.497 | 0.476 | 0.455 | 0.432 | 0.442 | 0.403 | 0.399 | 0.365 |
| Naive Bayes | **0.574** | **0.535** | **0.543** | **0.526** | **0.599** | **0.532** | **0.555** | **0.522** | 0.598 | 0.549 |

Table – S6: Accuracy across different ROIs for the classification shown by various models

| Model | No. of ROIs - 112 | No. of ROIs - 38 | No. of ROIs - 26 | No. of ROIs - 22 | No. of ROIs - 16 |
|---|---|---|---|---|---|
| K Neighbors Classifier | 0.427 | 0.487 | 0.416 | 0.478 | 0.375 |
| Random Forest Classifier | 0.433 | 0.477 | 0.366 | 0.491 | 0.326 |
| Logistic Regression | 0.512 | 0.468 | 0.388 | **0.538** | **0.649** |
| Decision Tree Classifier | 0.389 | 0.528 | 0.446 | 0.412 | 0.384 |
| Naive Bayes | **0.543** | **0.587** | **0.561** | 0.536 | 0.588 |

Table S7 represents the ROI pairs contributing highest towards the classification predicted by the algorithms when using 26 ROIs

Table – S7: Top 10 ROI Pairs contributing the classification using the dataset produced by 26 ROIs

| ROI Pair Codes | ROI - 1 | ROI - 2 |
|---|---|---|
| A146 | SMA1_R | iPreCG_R |
| A82 | ParOper_R | IFG3_R |
| A245 | SMG3_L | ParOper_R |
| A244 | SMG3_L | ParOper_L |
| A238 | SMG3_L | IFG5_L |
| A93 | pSTG_L | IFG1_R |
| A216 | SMG1_R | IFG4_R |
| A300 | tempMed_L | SMG3_R |
| A145 | SMA1_R | iPreCG_L |
| A178 | SMA2_R | IFG5_L |

Table S8 represents the ROI pairs contributing highest towards the classification predicted by the algorithms when using 22 ROIs

Table – S8: Top 10 ROI Pairs contributing the classification when used dataset produced by 22 ROIs

| ROI Pair Codes | ROI - 1 | ROI - 2 |
|---|---|---|
| A134 | SMA1_L | SMG3_R |
| A161 | SMA2_L | IFG5_R |
| A49 | SMG1_L | IFG3_R |
| A164 | SMA2_L | SMG1_L |
| A118 | tempMed_R | SMG3_L |
| A55 | SMG1_L | occMed_R |
| A160 | SMA2_L | IFG5_L |
| A168 | SMA2_L | tempMed_L |
| A133 | SMA1_L | SMG3_L |
| A141 | SMA1_R | IFG4_L |
